# Differential *in vivo* biodistribution of ^131^ I-labeled exosomes from diverse cellular origins and its implication in the theranostic application

**DOI:** 10.1101/566323

**Authors:** Mohammad H. Rashid, Thaiz F. Borin, Roxan Ara, Kartik Angara, Jingwen Cai, Bhagelu R. Achyut, Yutao Liu, Ali S. Arbab

## Abstract

Exosomes are critical mediators of intercellular crosstalk and regulator of cellular/tumor microenvironment. Exosomes have great prospects for clinical application as theranostic and prognostic probe. Nevertheless, the advancement of the exosomes research has been thwarted by limited knowledge elucidating the most efficient isolation method and their *in vivo* trafficking. Here we have showed that combination of two size-based methods using 0.20 µm syringe filter and 100k centrifuge membrane filter followed by ultracentrifugation method yields a greater number of uniform exosomes. We also demonstrated the visual representation and quantification of differential *in vivo* distribution of radioisotope ^131^I-labelled exosomes from diverse cellular origins, e.g., tumor cells with or without treatments (HET0016 and GW2580), myeloid-derived suppressor cells and endothelial progenitor cells. We also determined that the distribution was dependent on the protein/cytokine contents of the exosomes. The applied *in vivo* imaging modalities can be utilized to monitor disease progression, metastasis, and exosome-based targeted therapy.

**Abbreviations:**

bFGF: basic fibroblast growth factor
CSF1R: colony stimulating factor 1 receptor
CT: computed tomography
CTLA4: cytotoxic T-lymphocyte-associated protein 4
EGF: epidermal growth factor
EMT: epithelial to mesenchymal transition
EVs: extracellular vesicles
EPCs: endothelial progenitor cells
FasL: Fas ligand
G-CSF: granulocyte-colony stimulating factor
GM-CSF: granulocyte-macrophage colony-stimulating factor
HGF: hepatocyte growth factor
HSP: heat shock protein
ICAM-1: intercellular adhesion molecule 1
IFN-gamma: interferon gamma
IL – 1beta: interleukin-1 beta
IL – 1ra: interleukin-1 receptor antagonist
IL – 2: interleukin-2
IL – 4: interleukin-4
IL – 6: interleukin-6
IL – 7: interleukin-7
IL – 10: interleukin-10
IL – 12: interleukin-12
IL – 13: interleukin-13
IL – 17: interleukin-17
KC: keratinocyte-derived chemokine
LIX: lipopolysaccharide-induced CXC chemokine
M-CSF: macrophage colony-stimulating factor
MCP-1: monocyte chemoattractant protein 1
MDC: macrophage-derived chemokine
MDSCs: myeloid derived suppressor cells
MFP: mammary fat pad
MIP-1α: macrophage-inflammatory protein-1alpha
MMP-2: matrix metalloproteinase-2
MRI: magnetic resonance imaging
NIS: sodium iodide symporter
NTA: nanoparticle tracking analysis
PET: positron emission tomography
PF-4: platelet factor 4
RANTES: regulated on activation, normal T cell expressed and secreted
ROIs: region of interest
SDF-1α: stromal cell-derived factor-1
SEM: standard error of the mean
SPECT: single-photon emission computed tomography
SCF: stem cell factor
TAMs: tumor-associated macrophages
TEM: transmission electron microscopy
TIMP 2: tissue inhibitors of metalloproteinases 2
TLPC: thin layer paper chromatography
TME: tumor microenvironment
TNF-α: tumor necrosis factor-α
TSLP: thymic stromal lymphopoietin
UC: ultracentrifugation
VEGF-A: vascular endothelial growth factor A
VEGFR2: vascular endothelial growth factor receptor 2.

## Background

Exosomes are highly heterogeneous [1] endosomal origin lipid bilayered membranous vesicles (30–150 nm) produced by all cell types [2] and released into the biological fluids or cell culture medium. The initial concept of exosomes as “garbage bag” has been changed drastically as now exosomes are thought to be an integral part of intercellular communication. At any given time point, exosomes can contain all known bioactive constituents of a cell, including proteins, lipids and nucleic acid [3-7]. Besides their physiological functions such as maintenance of cellular-stemness, immunity [8, 9], tissue homeostasis, protein clearance [10], signaling [11, 12]; exosomes contribute to the pathophysiology of several diseases. The tumor cell-derived exosomes (TDEs) are capable of altering the tumor microenvironment (TME) by stimulating the secretion of growth factors and cytokines from TME associated cells. Thus TDEs play an imperative role in epithelial to mesenchymal transition (EMT) [3], immune escape [13], tumorigenesis, tumor growth, angiogenesis, invasion, cancer stemness, tumor drug resistance [14]. TDEs arbitrate tumor-associated immune suppression [15] and trigger pre-metastatic niche formation, acclimated to cancer-cell seeding prior the arrival of the first cancer cells [16]. Exosomes have immense prospects for clinical application as a diagnostic marker, monitoring treatment response and disease progression. Recently, exosome-based therapeutic delivery system is drawing tremendous attention owing to the distinctions between exosomes and synthetic nano-carriers. Nevertheless, few unsettled queries still affect the possible application of exosomes e.g., most efficient and reproducible approach for large-scale production of quality exosomes in a shorter time, the loading efficiency of therapeutic agents and precise detection of exosome biodistribution.

Despite the intense research for understanding the biological and pathophysiological functions of exosomes, only few studies have scrutinized exosome biodistribution. A breakthrough to investigate *in vivo* distribution and track the exosomes is immensely desired for safe and effective clinical application. Yet, to track their whereabouts, only few effective methods are reported that mostly adopted fluorescent and bioluminescence imaging either labeling them with lipophilic membrane dye or manipulating them to exhibit a membrane reporter. Contrarily, the utilization of nuclear medicine imaging techniques, such as single-photon emission computed tomography (SPECT) or positron emission tomography (PET) that are non-invasive imaging, can be combined with anatomical imaging such as, computed tomography (CT) or magnetic resonance imaging (MRI) for exosome localization can be barely found in the published articles. These nuclear imaging techniques have indisputable advantages over fluorescent and bioluminescence imaging, owing to their excellent sensitivity for the deeper tissues and quantitative measurement potential of the clinical grade labeling radioisotopes (^99m^Tc, ^131^I, ^111^In-oxine).

Here, we proposed a simpler and quicker isolation technique by a combination of size based and ultracentrifugation method. We also demonstrated the visual representation and *in vivo* distribution of exosomes, isolated from tumor cells with or without treatment; myeloid-derived suppressor cells (MDSCs) and endothelial progenitor cells (EPCs) in metastatic breast cancer animal models by SPECT/CT. We also showed that differential biodistribution is related to the protein/cytokine contents of the collected exosomes.

## Methods

Description of cell lines, nanoparticle tracking analysis, flow cytometry, isolation of MDSCs and EPCs, protein quantification, thin layer paper chromatography, quantitative analysis of radioactivity, *ex vivo* gamma activity and statistical analysis are described in Supplementary Material.

### Ethics statement

All the experiments were performed according to the National Institutes of Health (NIH) guidelines and regulations. The Institutional Animal Care and Use Committee (IACUC) of Augusta University (protocol #2014–0625) approved all the experimental procedures. All animals were kept under regular barrier conditions at room temperature with exposure to light for 12 hours and dark for 12 hours. Food and water were offered ad libitum. All efforts were made to ameliorate the suffering of animals. CO2 with secondary method was used to euthanize animals for tissue collection.

### Exosome isolation

Exosomes were isolated from the culture supernatant of 4T1 and AT3 tumor cell lines. Briefly, 5×10^6^ tumor cells were plated in 175cm^2^ flasks and grown overnight with 10% FBS complete media in normoxia (20% oxygen). The media was removed and replenished by exosome free complete media. Exosomes were depleted from the complete media by ultracentrifugation for 70 minutes at 100,000x g using a Ultracentrifuge (Beckman Coulter) and SW28 swinging-bucket rotor. Then the cells were treated with control (DMSO), colony stimulating factor-1 receptor (CSF1R) antagonist (GW2580, 1µM) and 20-HETE synthase inhibitor (HET0016, 100µM) in hypoxia (1% oxygen) for 48 hours. The cell culture supernatant was centrifuged at 1500 rpm for 10 minutes to get rid of cell debris. We employed five different methods as follows – 1) ultracentrifugation only by initial step with 10,000x g for 30 minutes followed by two steps with 100,000x g for 70 minutes each, 2) size-based method by passing through 0.20 µm syringe filter (Corning, USA) followed by centrifugation with 100k membrane tube (Pall Corporation, USA) at 3900 rpm for 30 minutes, 3) Combination of two steps of size-based method by passing through 0.20 µm syringe filter and centrifugation with 100k membrane tube at 3900 rpm for 30 minutes followed by single step of ultracentrifugation at 100,000x g for 70 minutes, 4) combination of one step size-based method by passing through 0.20 µm and single ultracentrifugation at 100,000x g for 70 minutes and 5) commercially available density gradient separation by total exosome isolation reagent (Invitrogen™, USA). The reagent was added to the culture supernatant sample and incubated overnight at 4° C. The precipitated exosomes were recovered by centrifugation at 10,000x g for 60 minutes.

### Tumor model

Both 4T1 and AT3 cells expressing the luciferase gene were orthotopically implanted in syngeneic BALB/c and C57BL/J6 mice, respectively (Jackson Laboratory, USA). All the mice were between 5-6 weeks of age and weighing 18-20g. Animals were anesthetized using a mixture of Xylazine (20mg/Kg) and Ketamine (100 mg/Kg). Either 50,000 4T1 cells or 100,000 AT3 cells in 50µL matrigel (Corning, USA) were injected into the right mammary fat pad.

### Radiolabeling of exosomes using Iodine-131 (^131^I)

Isolated TDEs were labeled by Pierce™ Iodination Beads (Thermo Scientific™). In short, 4-5 iodination beads were cleaned with sterile normal saline and allowed to air dry. The beads were then added directly to 5 mCi of ^131^I solution (Cardinal Health, Inc.) and then incubated at room temperature. After 5 minutes, exosomes resuspended in PBS were added to the reaction tube and incubated at room temperature for 30 minutes. To stop the iodination reaction, the beads were taken out from the reaction tube. To get rid of free ^131^I, the labeled exosomes were washed and centrifuged with extra PBS using a 100k membrane tube at 3900 rpm for 10 minutes.

### *In vivo* SPECT/CT imaging

After the intravenous injection of 350±50 µCi of ^131^I-labeled exosomes in 100µL into the tail vein, all animals underwent SPECT-CT scanning. During the whole procedure, the animals were anesthetized and maintained using a combination of 1.5% isoflurane and 1 L/min medical oxygen flow and their body was immobilized in an imaging chamber to restrain movements. Whole body CT followed by SPECT-imaging was acquired by a nanoScan 4-headed micro-SPECT-CT scanner (Mediso, USA). The image acquisitions were commenced 3 hours after the injection of ^131^I-labeled exosomes. The reconstructed image size was 205×205×205µm.

### Protein array

Exosomal proteins were evaluated for the expression profiles of 44 factors in duplicate by mouse cytokine antibody array (RayBiotech, Inc.). 500μg of protein sample was loaded to the membrane and the chemiluminescent reaction was detected by using LAS-3000 imaging machine (Fuji Film, Japan). All signals emitted from the membrane were normalized to the average of 6 positive control spots of the corresponding membrane using ImageJ software.

## Results

### Optimizing the exosome isolation method

To optimize exosome isolation method, we collected exosomes from 4T1 cell culture supernatant using five different techniques-1) ultracentrifugation (UC) only, 2) size based only (0.20 µm and 100k membrane), 3) combination of size based (0.20 µm and 100k membrane) and UC, 4) combination of single size based (0.20 µm) and UC and 5) density gradient separation. Among the different methods employed, density gradient separation technique (#5) yielded most concentrated exosomes with 1.1×10^11^ particles/mL, while (#1) UC alone and its (#4) combination with size based (0.20 µm) yielded the lowest concentration of 1.8×10^10^ particles/mL and 1.7×10^10^ particles/mL, respectively (**Figure 1A**). Combination of size based (0.20 µm and 100k) followed by UC (#3) yielded at a concentration of 4.2×10^10^ particles/mL. However, after the isolation process, there was visible sedimentation of co-isolated impurities and polymer or reagent along with the exosomes isolated by density gradient separation. Mean diameter of exosomes isolated by (#1) UC alone was 149.7±64.3 nm, (#3) combination of size based (0.20 µm and 100k membrane) and UC was 126.7±52.9 nm and (#5) density gradient method was 135.1±70.8 nm (**Figure 1B**). Size distribution curve of density gradient separation showed a wider base with a thick tail extending towards smaller size (**Figure 1C**). We also examined the common exosomal markers, CD9 and CD63, in all the samples by flow cytometry (**Figure 1D**). There was no significant difference between density gradient separation and the combination of size based and UC technique. Transmission electron microscopy (TEM) images for the exosomes isolated by a combination of size based and UC method showed normal morphology of exosomes without any distortion (**Figure 1E**).

**Figure 1.**
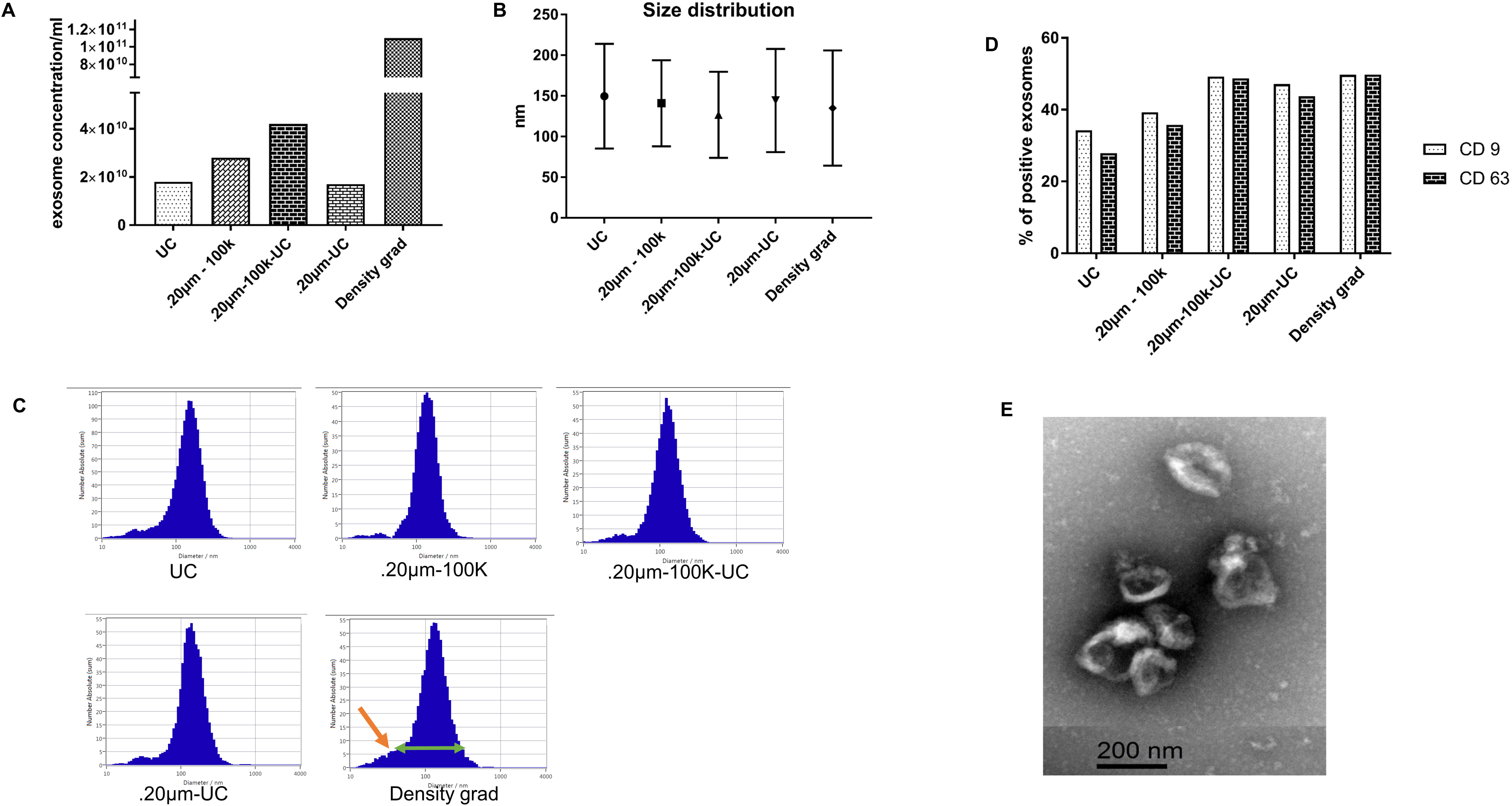
**(A)** NTA analysis showing exosomes concentration per mL and **(B and C)** size distribution from five different isolation methods. **(D)** Flow-cytometric analysis of common exosome markers (CD9 and CD63). **(E)** Transmission electron microscopy (TEM) image for the exosomes isolated by (#3) combination of size based and UC method, scale bar depicts 200 nm.

### Binding efficiency and serum stability of of ^131^I-labeled exosomes

For confirming the binding efficacy of ^131^I to the exosomes and serum stability, we implemented thin layer paper chromatography (TLPC) [17]. Most of the free ^131^I alone moved from the spotted point in the bottom to the top half (**Figure 2A**). Contrarily, ^131^I bound to the exosomes remained at the bottom and barely moved to the top half, indicating ^131^I bound to the exosomes and very little dissociation of the ^131^I from the exosomes (**Figure 2B**). We appraised the labeling stability of ^131^I-labeled exosomes in serum by incubating ^131^I labeled exosomes with 20% FBS for 4 and 24 hours in 37° C followed by TLPC similar to the previous experiment. Identical to the ^131^I bound exosomes, ^131^I from serum-challenged ^131^I labeled exosomes hardly moved to the top, implying the labeling of exosomes with ^131^I was stable in serum even after 24 hours (**Figure 2C**).

**Figure 2.**
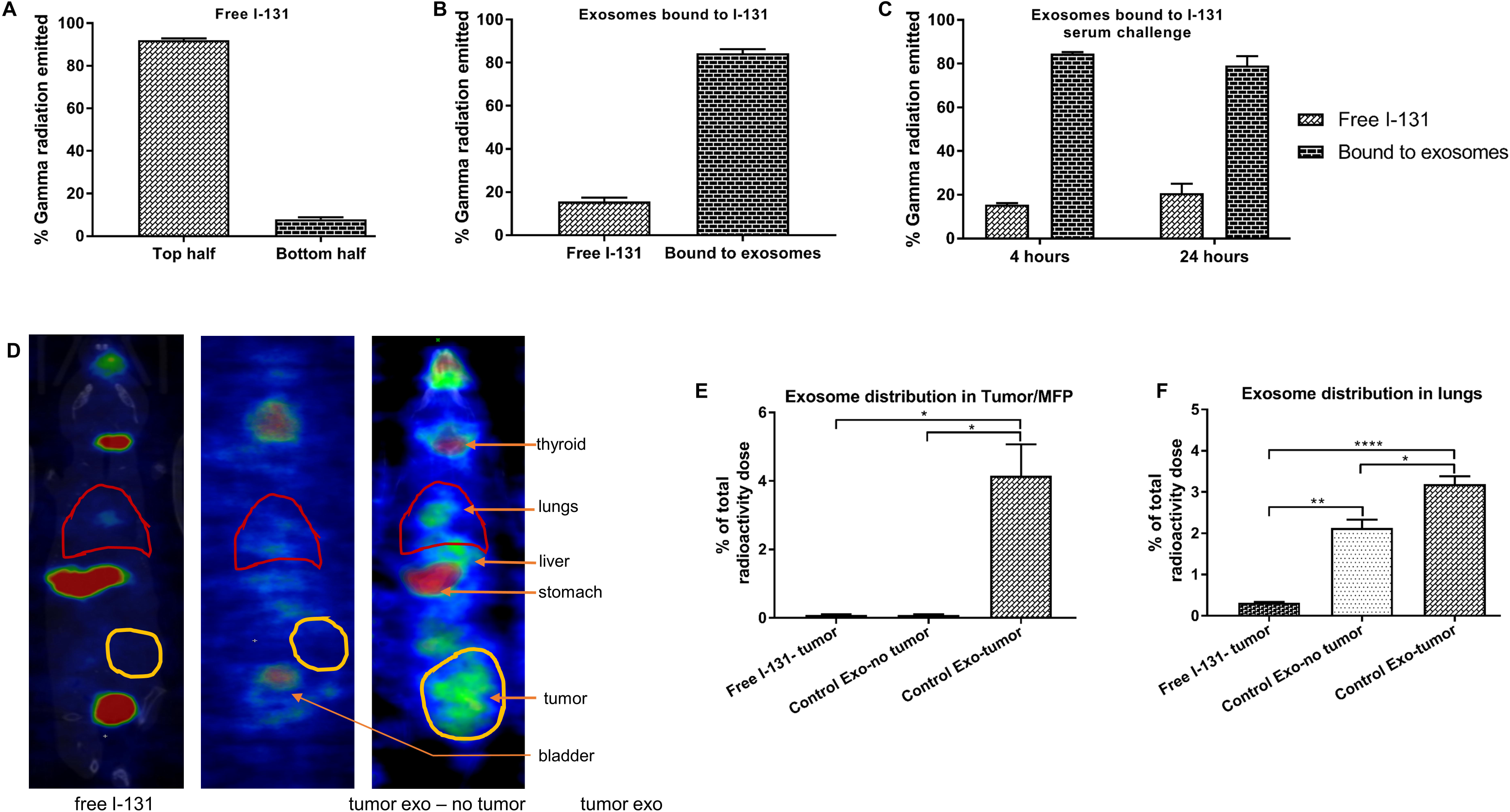
Binding of ^131^I to the exosomes and serum stability was confirmed by thin layer paper chromatography (TLPC). **(A)** Major proportion of the free ^131^I alone moved from the spotted point in the bottom to the top half, confirming the efficacy of the eluent. **(B)** Binding of ^131^I to the exosomes was validated as very less percentage of dissociated free ^131^I from the labeled exosomes moved to the top half. **(C)** Serum stability of ^131^I bound exosomes was very high as very small amount of free ^131^I disengaged from the bound exosomes to move to the top half. **(D)** Reconstructed and co-registered *in vivo* SPECT/CT images (coronal view) at 3 hrs, from the animals injected with ^131^I-labeled tumor exosomes. (**E and F**) Free I-131 and exosome distribution in the tumors and lungs. Quantitative data are expressed in mean ± SEM. *P<.05, ***P<.001. n = 3.

### Detection and quantification of radioisotope ^131^I-labeled tumor cell-derived exosomes in the primary tumor and metastatic site

^131^I labeled-TDEs in 100 µL solution was injected into tumor-bearing (tumor exo) and tumor-free (tumor-exo – no tumor) animals via tail-vein. After 3, 24 and 48 hrs of administration, the animals were scanned by SPECT/CT, and the reconstructed images were analyzed by ImageJ. We also injected free ^131^I in tumor-bearing mice (free I-131) to determine the uptake of free ^131^I to the tumors. We observed an ample amount of radioactive intensity after 3 hrs at the primary tumor site and metastatic site (lung) in the animals that received ^131^I-labeled TDEs (**Figure 2D**). There was almost no radioactivity in the tumor area, and negligible radioactivity in the lungs of the group injected with free ^131^I. The group of animals without any tumor and injected with radiolabeled TDEs showed almost no radioactivity in the mammary fat pad but plenty of radioactivity in the lungs (**Figure 2E and 2F**). We observed notable radioactivity in thyroid and stomach. Radioactivity was also high in the bladder for the renal clearance.

### Distribution of MDSCs, EPCs and HEK293-derived exosomes in primary and metastatic site

Next, we wanted to see the biodistribution of exosomes derived from other cell types that play crucial role in tumor progression and metastasis. As a non-cancerous, non-specific cell line, we used human embryonic kidney 293 (HEK293) cells. MDSCs were collected from the spleen of tumor-bearing mice with more than 99% purity, and EPCs were isolated from the bone marrow of normal mice with more than 85% purity (**Figure 3A**). TDEs (tumor exo) were collected from the cell culture supernatant. Mean diameter of isolated exosomes from HEK293 cells (HEK293 exo), MDSCs (MDSC exo) and EPCs (EPC exo) was 97.6 nm, 131.1 nm, and 140.1 nm respectively (**Figure 3B and 3C**). Three hours after injecting the ^131^I-labeled exosomes intravenously in tumor-bearing animals, exosomes from all groups accumulated in the primary breast tumor and metastatic site in the lungs except the HEK293 exo (**Figure 3D**). Interestingly, exosomes from the EPCs were abundantly located in the primary tumor site, and ample amount of exosomes from the MDSCs were visualized more in the metastatic site-lungs than any other groups (**Figure 3E and 3F**).

**Figure 3.**
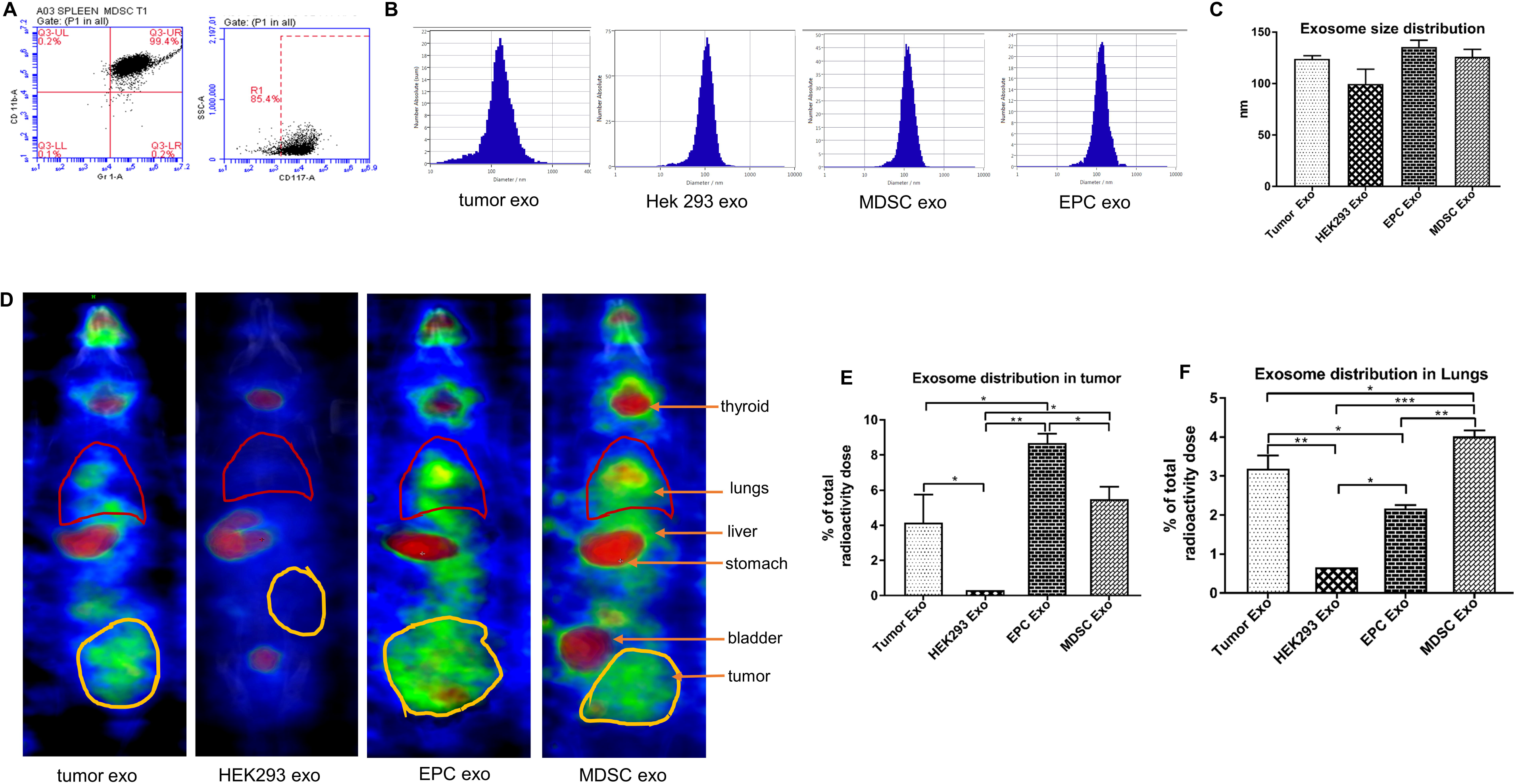
**(A)** Flow cytometric analysis of isolated MDSCs, showing more than 99% of cells are positive for CD11b+ and Gr1, and EPCs showing more than 85% positive for CD117. **(B and C)** NTA is showing size distribution curve and mean size of the exosomes from tumor cells (tumor exo), HEK293 cells (HEK293 exo), myeloid-derived suppressor cells (MDSC exo) and endothelial progenitor cells (EPC exo). **(D)** Reconstructed and co-registered *in vivo* SPECT/CT images (coronal view) of the abovementioned ^131^I labeled exosomes after 3 hrs of intravenous injection in tumor-bearing mice. **(E)** Quantification of radioactivity in tumor showed significant aggregation of EPC exo contrast to control exo and MDSC exo. **(F)** Quantification of radioactivity in lungs showed highest percentage of radioactivity in mice injected with MDSC exo. There was no retention of non-specific HEK293 exo in either tumor or lungs. Quantitative data are expressed in mean ± SEM. *P<.05, ***P<.001. n = 3.

### *Ex-vivo* gamma activity measurement of a different organ

After the final scan, we euthanized the animals, measured the weight and emitted gamma activity of individual harvested organs. While most of the organs as a whole showed negligible radioactivity, only the tumor and the liver retained a significant load of radioactivity in all the groups (**Supplemental Figure S1A**). Interestingly, lungs from the animals injected with MDSC-exo showed considerably higher radioactivity than the other groups. For the EPC-exo, the primary tumor showed notably higher gamma activity than the other organs. We also calculated the radioactivity per milligram of the weight of an individual organ that demonstrated similar changes of radioactivity as of the whole organ. The non-tumor bearing group, injected with ^131^I labeled tumor exosomes showed gamma activity mostly in the liver (**Supplemental Figure S1B**).

### Cytokine array of MDSCs and EPCs-derived exosomal protein content

To understand variation in biodistribution, we analyzed exosomal protein contents from MDSCs and EPCs compared with those of TDEs by membrane-based cytokine array. Angiogenic factors such as Endoglin, Intercellular adhesion molecule-1 (ICAM-1) and Platelet factor-4 (PF-4) were increased in both MDSC exo and EPC exo (**Figure 4**). The level of basic fibroblast growth factor (bFGF) and vascular endothelial growth factor receptor-2 (VEGFR2) was high in MDSC exo. Epidermal growth factor (EGF) was high in both MDSC and EPC exo than the tumor exo. Among the invasion factors, E-cadherin and E-selectin level was increased in both MDSC and EPC exo. Notably, matrix metalloproteinase-2 (MMP-2) was increased in MDSC exo compared to tumor and EPC exo. Factors for myeloid activation and function such as granulocyte-colony stimulating factor (G-CSF), granulocyte-macrophage colony stimulating factor (GM-CSF), macrophage-colony stimulating factor (M-CSF) and monocyte chemoattractant protein-1 (MCP-1/CCL2) were over-expressed in both MDSC and EPC exo than the tumor exo. In addition, MDSC exo also had higher expression of macrophage-inflammatory protein-1-alpha (MIP-1α/CCL3) and stromal cell-derived factor-1 (SDF-1α). T-cell function modulating factors like interleukin (IL)-2, IL-7, L-selectin, thymic stromal lymphopoietin (TSLP) were increased in MDSC and EPC exo. From the immunomodulatory cytokines, IL-2, IL-4, keratinocyte-derived chemokine (CXCL1/KC), C-X-C motif chemokine-5 (CXCL5/LIX), chemokine (C-C motif) ligand-5 (CCL5/RANTES) and tumor necrosis factor-α (TNF-α) were significantly high in both MDSC and EPC exo than the control exo. Furthermore, there was an obvious increase of IL-13 and IL-1 receptor antagonist (IL-1ra) in MDSC exo.

**Figure 4.**
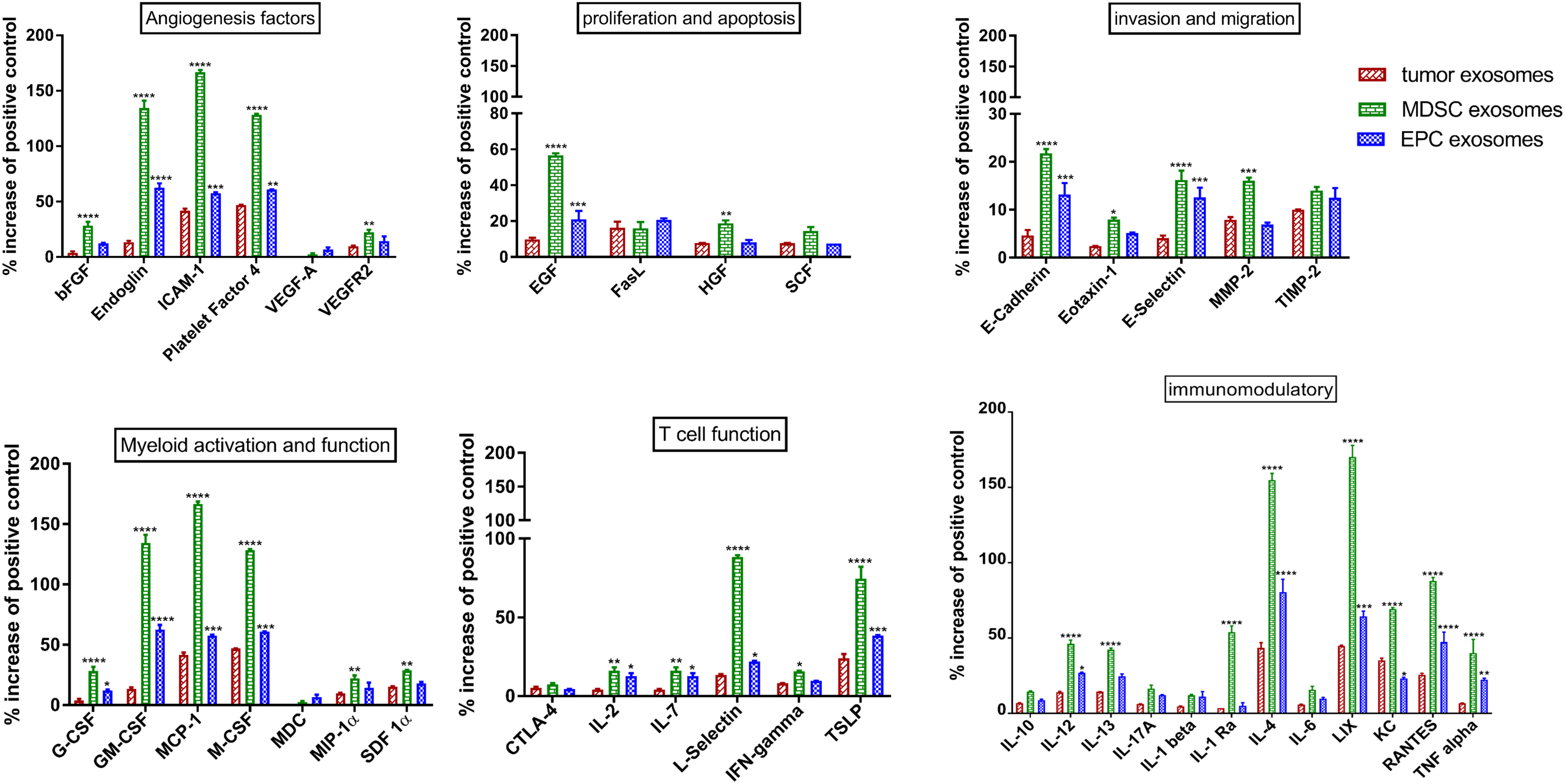
*In vitro* quantification of the level of cytokines by membrane-based array in the protein samples, collected from the exosomes of AT3 tumor cells, MDSCs, and EPCs. Exosomal proteins from MDSCs and EPCs showed significant increase in some pro-angiogenic factors (Endoglin, ICAM-1, PF-4), immune modulatory factors (TNF-α, IL-4, IL-12, LIX, RANTES), myeloid activation and function factors (GM-CSF, G-CSF, M-CSF and MCP-1), T cell-related factors (IL-2, IL-7, L-selectin, TSLP) than the exosomal protein from tumor cells. There was significant upregulation of bFGF, VEGFR2, EGF, HGF, MMP-2, MIP-1α, SDF-1α, IFN-gamma, IL-13 and IL-1ra level in MDSCs-derived exosomes than EPCs and tumor-derived exosomes. Quantitative data are expressed in mean ± SEM. *P<.05, **P<.01, ***P<.001, ****P<.0001. n = 2.

### Distribution of exosomes from treated tumor cells

To explore further, we wanted to investigate whether exosomes collected from tumor cells treated with drugs that decrease cancer growth and metastasis, would distribute differently *in vivo*. Previously our group showed that HET0016 and GW2580 treatment decreases lung metastasis of breast tumor-bearing mice [18, 19] and HET0016 decreases neovascularization, tumor growth with increased survival in glioblastoma model [18, 19]. GW2580, a selective small molecule kinase inhibitor of CSF1R and HET0016 is a highly selective inhibitor of CYP4A in the arachidonic acid pathway, that have been shown to decrease recruitment of tumor-infiltrating myeloid cells and decreased tumor-associated macrophages (TAMs) polarization towards M2 macrophages [20]. There was no significant differences in exosome size and marker (CD9 and CD63) after GW2580 and HET0016 treatment of cells (**Figure 5A**, **5B and 5C**). After labeling with ^131^I, the exosomes were injected in tumor-bearing mice and in *vivo* imaging was acquired by SPECT/CT after 3 hours, 24 hours and 48 hours. There was increased radioactivity at the primary tumor site in all the groups (**Figure 5D**). Though there was an increased localization of the HET exo in the primary tumor, it was not statistically significant (**Figure 5E**). However, there was an ascertainable decline of radioactivity in the lungs of GW exo, and HET exo groups compared to that of control exo groups (**Figure 5F**).

**Figure 5.**
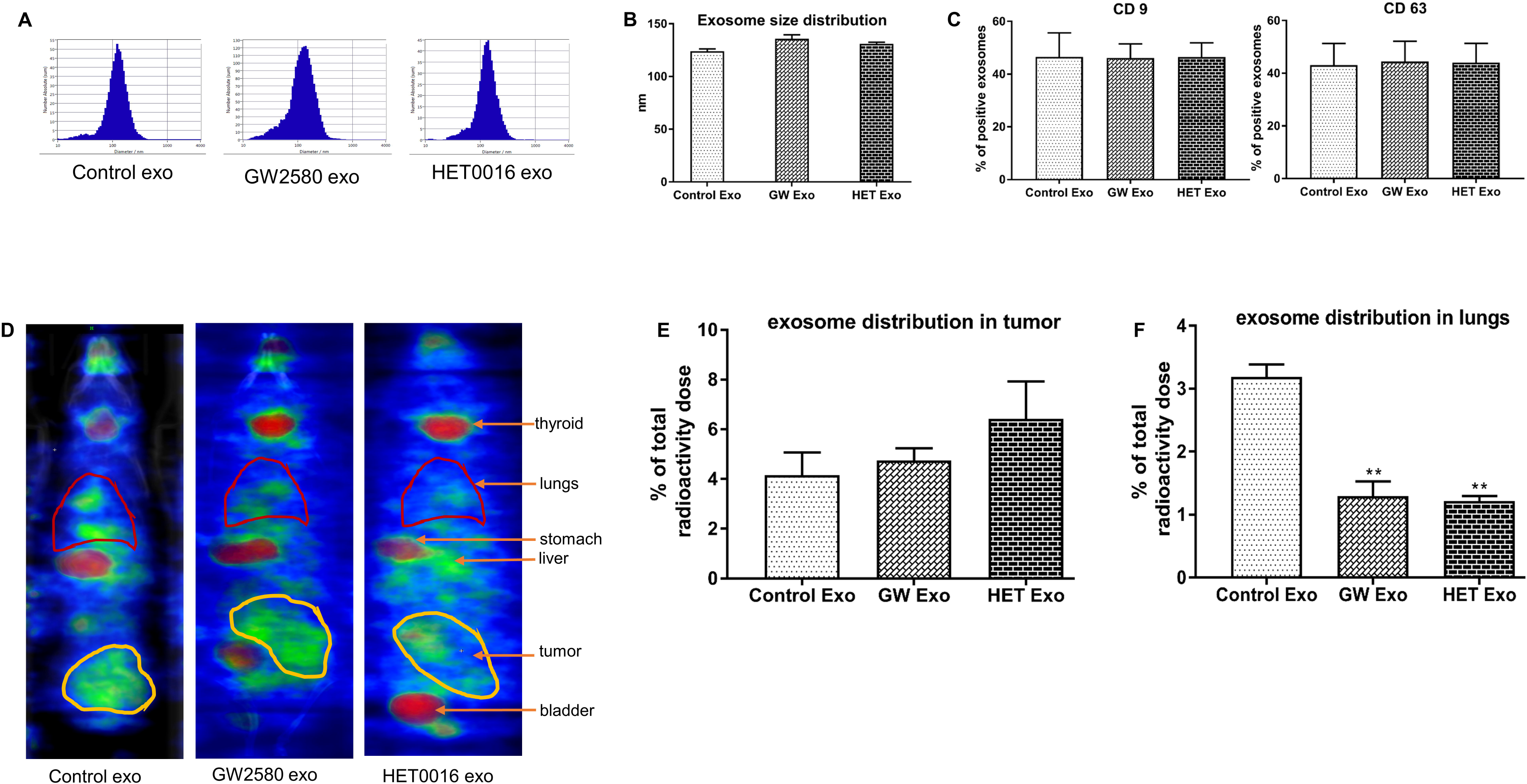
**(A and B)** NTA analysis showing no significant differences in the size distribution of exosomes isolated from tumor cells (control exo), GW2580 treated tumor cells (GW2580-exo), and HET0016 treated tumor cells (HET0016-exo). **(C)** Flow-cytometric analysis of exosomal markers-CD9 and CD63 for abovementioned treatment conditions. **(D)** Reconstructed and co-registered *in vivo* SPECT/CT images (coronal view) of the animals injected with abovementioned ^131^I labeled exosomes after 3 hrs. **(E)** Quantification of radioactivity in tumor showed insignificant higher aggregation in mice injected with 131I-labeled GW2580 exo and HET0016 exo compared to control exo. **(F)** Quantification of radioactivity in lungs showed significant reduction of exosomes localization in GW 2580 exo and HET0016 exo than the control exo injected groups. Quantitative data are expressed in mean ± SEM. *P<.05, **P<.01, ***P<.001, ****P<.0001. n = 3.

### Distribution in other organs and clearance

In addition to the tumor and lungs, we also measured the radioactivity in the other organs and clearance over time (**Supplemental Figure S2**). After 3 hours, the highest percentage of radioactivity was observed in the urinary bladder area that was subsequently cleared off. Considerable amount of radioactivity was noticed in the stomach, which was scarcely detectable in subsequent scanning. Among the organs, liver, lungs, and tumor retained the perceptible amount of radioactivity over the time. Furthermore, thyroid gland in all the groups showed no substantial change of radioactivity in all three scans (3, 24 and 48 hrs), suggesting no dissociation of the ^131^I from the exosomes that may eventually increase the free ^131^I uptake by the thyroid.

### Cytokine array of tumor-derived exosomal protein content after treatment

Finally, because of the differential biodistribution of exosomes isolated from tumor cells with or without treatment, we wanted to see if the treatment of 4T1 tumor cells with GW2580 and HET0016 affects the protein contents of exosomes. We extracted the exosomal proteins from the 4T1 cells without any treatment (control), GW2580 treated cells, and HET0016 treated cells. We performed cytokine array analysis to evaluate the changes of 44 cancer-related factors for these three samples. Among the angiogenesis factors, VEGFR2, ICAM-1, bFGF were significantly reduced in exosomes from the treated cells than the control cells (**Figure 6**). From the chemotactic factors, level of KC and macrophage-derived chemokine (MDC) was significantly downregulated in GW2580 exo and HET0016 exo respectively. Among the immune-modulatory cytokines, IL-6, TNF-α, IL-12, IL-10, IL-13 were decreased, and IL-4 was increased in the treated samples. Level of few invasive factors-E-cadherin, Eotaxin, E-selectin and tissue inhibitors of metalloproteinases-2 (TIMP-2) was significantly reduced in exosomes from GW2580 treated cells. Interestingly, both GW2580 and HET0016 treatment of tumor cells increased the Fas ligand (FasL/CD95L) in the exosomes.

**Figure 6.**
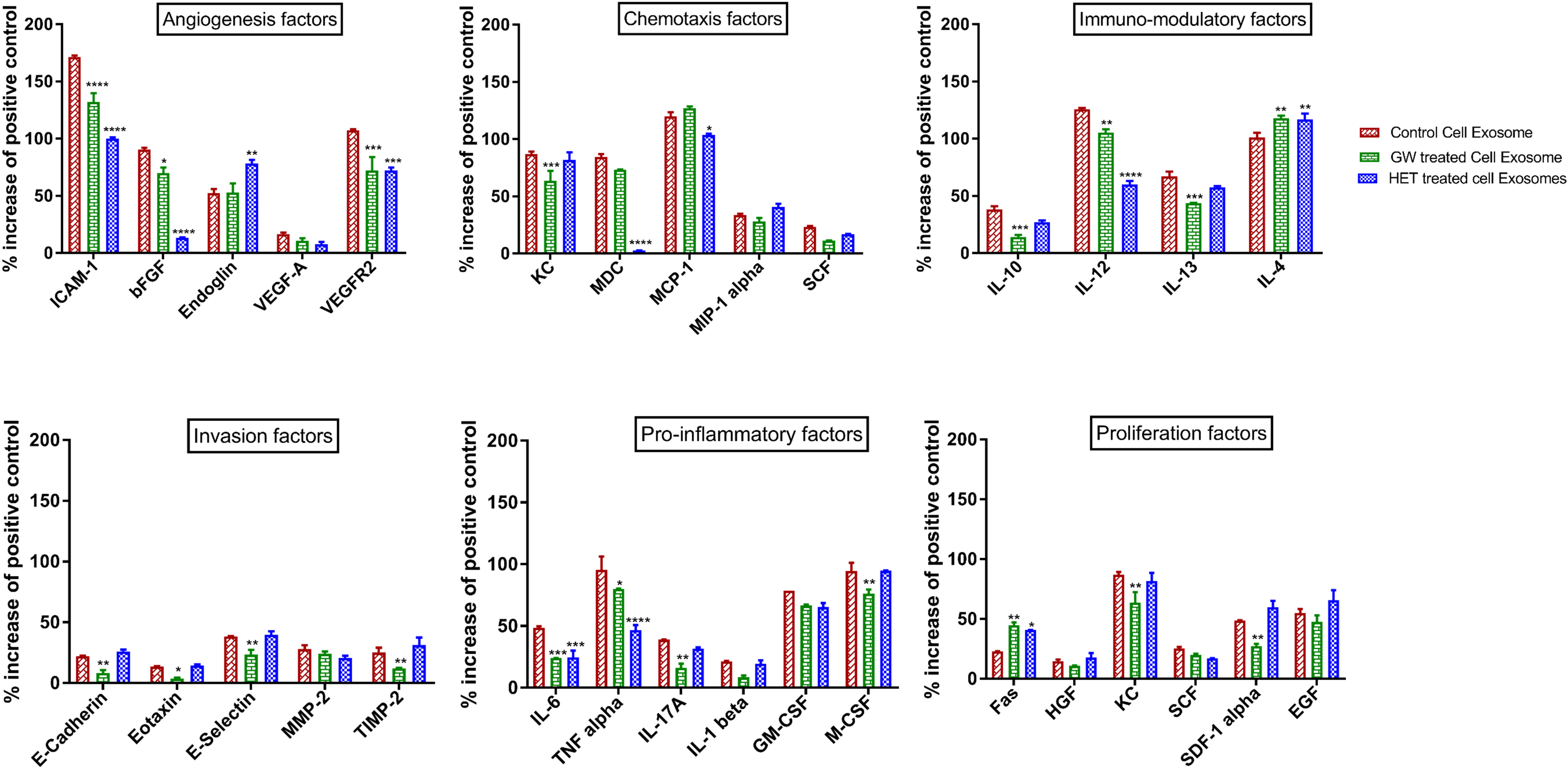
*In vitro* quantification of the level of cytokines by membrane-based array in the protein samples, collected from the exosomes of 4T1 tumor cells without any treatment and with GW2580 and HET0016 treatment for 48 hrs.Exosomal proteins from treated cells showed significant decrease of some pro-angiogenic (VEGFR2, ICAM-1, bFGF), immune modulatory (TNF-α, IL-6, IL-12, IL-10, IL-13) and chemotactic (KC, MDC) factors. FasL level was increased in both exosomal protein samples from treated cells. Quantitative data are expressed in mean ± SEM. *P<.05, **P<.01, ***P<.001, ****P<.0001. n = 2.

## Discussion

To expedite the research and clinical application of the exosomes, it is fundamental that exosomes are explicitly and more effectively isolated from a wide spectrum of impurities e.g., virus, proteins, apoptotic bodies. Until now, mainly five isolation techniques have been developed by exploiting a particular trait, such as their size, density and surface markers [21]. Here, we compared five different isolation techniques and implemented a modified reproducible protocol of isolating quality exosomes more efficiently for downstream experiments in a shorter time. UC only method takes approximately 4-5 hrs to isolate exosomes from the culture supernatant, and density gradient-based technique needs long incubation (overnight) time. Although size-based methods (alone or in combination) need shortest time, it is not possible to pellet down the exosomes from the media and subsequent wash. Our optimized method by a combination of size based using 0.20 µm syringe filter followed by 100k centrifuge membrane filter and ultracentrifugation method requires less than 2 hrs, and with the NTA we demonstrated that it yields greater amount of exosomes with uniform size. Also, exosomes isolated by our optimized method showed a comparable percentage of common exosome markers, CD9 and CD63. Using 0.20 µm syringe filter allows passing of vesicle size <200 nm, ensuring the exclusion of other larger extracellular vesicle types. Following 100k membrane centrifugation makes sure of getting rid of most of the protein impurities that could be co-isolated with the exosomes by other methods. Although, density gradient separation techniques yielded more exosomes, the size distribution curve was not uniform, and there were visible impurities along with isolated exosomes. We did not compare the immune-affinity capture based isolation method as it separates only the specific subgroup of exosomes that possesses the antigen of interest.

A few articles have reported the tumor targeting and metastatic tumor behavior of exogenously administered exosomes. While most of them adopted either fluorescence imaging [22-26] or bioluminescence imaging [27], only one article reported limited tumor accumulation of injected exosomes (labeled with ^111^In-oxine) by nuclear imaging [25]. However, the authors did not demonstrate visualization in tumor and metastatic site *in vivo.* Fluorescence imaging shows inferior tissue penetration, auto-fluorescence of biological tissues, low resolution and often requires euthanasia. Bioluminescence has low sensitivity for deeper organs with the necessity of injecting toxic substrates to generate optical signals and incompatible for sequential imaging [28]. We administered ^131^I-labeled TDEs followed by CT and SPECT imaging at three different time points. Reconstructed and co-registered *in vivo* SPECT/CT images of 3hrs post-injection showed significant accumulation of exosomes in primary tumor in tumor-bearing mice compared to the free ^131^I injected group and mammary fat pad (MFP) of non-tumor bearing mice. Noteworthy to mention that, there was almost no accumulation of TDEs in normal MFP, but they accumulated even in the normal lungs, implying the propensity of the breast TDEs for the future metastatic site. Thyroid and stomach showed significant radioactivity due to the presence of sodium-iodide symporter (NIS), which is an intrinsic plasma membrane glycoprotein, actively arbitrates iodide transport into the thyroid follicular cells [29] and several extra-thyroidal tissues, e.g., salivary glands, gastric mucosa and lactating mammary glands [29, 30]. However, the NIS expression level is lower in extra-thyroidal tissues than in thyroid tissues, and longstanding retention of iodide does not occur in these tissues [31]. That is why initial high radioactivity in the stomach was almost undetectable in subsequent scanning.

MDSCs and EPCs are critical components of TME, have differential functions in respect to tumor growth and metastasis. MDSCs are a heterogeneous population of immature myeloid cells that are directly implicated in the escalation of tumor metastases by partaking in the EMT, tumor cell invasion, promoting angiogenesis and formation of pre-metastatic niche [32, 33]. EPCs are progenitor cells with the ability to differentiate into endothelial cells and play a pivotal role in tumor growth and metastasis by tumor neovascularization, promoting the transition from micro to macro metastasis. Even though critical roles of MDSCs and EPCs in tumor progression and metastasis have been studied extensively, the similar roles of exosomes derived from these cells are yet to be explored. Only few articles are available on exosomes derived from MDSCs [14, 34-37] and EPCs [38-40], but not directly related to their contribution in cancer progression and metastasis. Interestingly, we observed the highest retention of MDSCs derived exosomes in lungs of breast tumor-bearing animals than any other groups, highlighting their metastatic properties.

On the other hand, we noticed a maximal load of accumulation of EPCs derived exosomes in the primary tumor than any other groups, which could be due to their neovascularization effects in the TME. *Ex vivo* measurements of radioactivity from individual organs by gamma counter also correspond to the *in vivo* analysis. To substantiate prior findings, we investigated the level of tumor-associated cytokines in exosomal protein content from MDSCs and EPCs compared to the tumor cells. There was a significant increase of endoglin, ICAM-1 and PF-4 level in the MDSC exo and EPC exo than the tumor exo. MDSC exo also showed higher expression of bFGF and VEGFR2. Endoglin [41], VEGFR2 [42], ICAM-1 [43] and bFGF [44] are established factors that promote neovascularization. Myeloid activation and function relevant cytokines such as GM-CSF, G-CSF, M-CSF, and MCP-1 were over-expressed in exosomal content from both MDSCs and EPCs.

Furthermore, MDSCs exo contained a higher level of MIP-1α and SDF-1α. GM-CSF and G-CSF potently induce the expansion of MDSCs leading to enhanced metastatic growth [45]. Not to mention, GM-CSF also promotes tumor proliferation and invasion while M-CSF instigates tumor invasion [46]. TSLP and IL-7 were over-expressed in MDSC and EPC exo, that drive the differentiation of Th2 cells followed by secretion of IL-13 and IL-4; leading to the recruitment and activation of MDSCs and TAMs, which promote metastasis by secreting tumor cell migration-stimulating factors such as SDF-1α, IL-6, MMP – 2, LIX, KC, RANTES and TNF-α [47, 48]. All the cytokines were significantly more in MDSCs-derived exosomes than EPCs exo or TDEs suggesting their prospective role in tumor proliferation, angiogenesis, invasion, and metastasis. While we used non-cancerous cells (HEK293)-derived exosomes, we did not see any retention of exosomes in primary tumor or metastatic site indicating that only the exosomes from tumor cells or TME-associated cells have the propensity to be distributed in tumor or metastatic area.

We explored the possibility that, chemotherapeutics or cell-targeted therapies may change the behavior and contents of the TDEs. We isolated exosomes from tumor cells with (GW2580, HET0016) or without treatment (control). Following intravenous injection of ^131^I-labeled exosomes, we observed more accumulation of treated exosomes in the tumor area, albeit it was not statistically significant. We also observed a significant reduction of radioactivity in lungs of the animal groups injected with GW2580 exo and HET0016 exo compared to the group injected with control-exo, elucidating the diminution of the metastatic competency of the exosomes after treating the parent cells with the therapeutics. For validating this finding, we profiled 44 cancer-related factors in the exosomal contents by cytokine array. Distinguishable cancer-promoting factors that declined following the treatment of cancer cells than the control are - VEGFR2, ICAM-1, bFGF, KC, TNF-α, IL-6, IL-12, IL-10, IL-13. KC promotes the growth of primary tumor and formation of a pre-metastatic niche in different tumor model [49, 50]. Elevated levels of IL-6 are associated with aggressive tumor growth, poor response to therapies, poor prognosis and shorter survival [51]. It should be noted that FasL level was elevated in the exosomes from treated cancer cells. Up-regulation of FasL often occurs following chemotherapy, from which the tumor cells attain apoptosis resistance [52, 53].

In summary, we favorably demonstrated a simple, rapid, high yielding exosome isolation technique utilizing a combination of size based and ultracentrifugation method. As per our knowledge, we are the first group to demonstrate the exact visualization and quantification of radioisotope-labeled exosomes from different cell types with or without treatment, in the primary solid tumors and metastatic area, which are dependent on the protein/cytokine contents of the exosomes. Our imaging technique and quantification of exosomes could be applied for potential metastatic site prediction, monitoring tumor progression and targeting efficacy of exosome-based therapy, thus unlocking a theranostic potential for these exosomes.

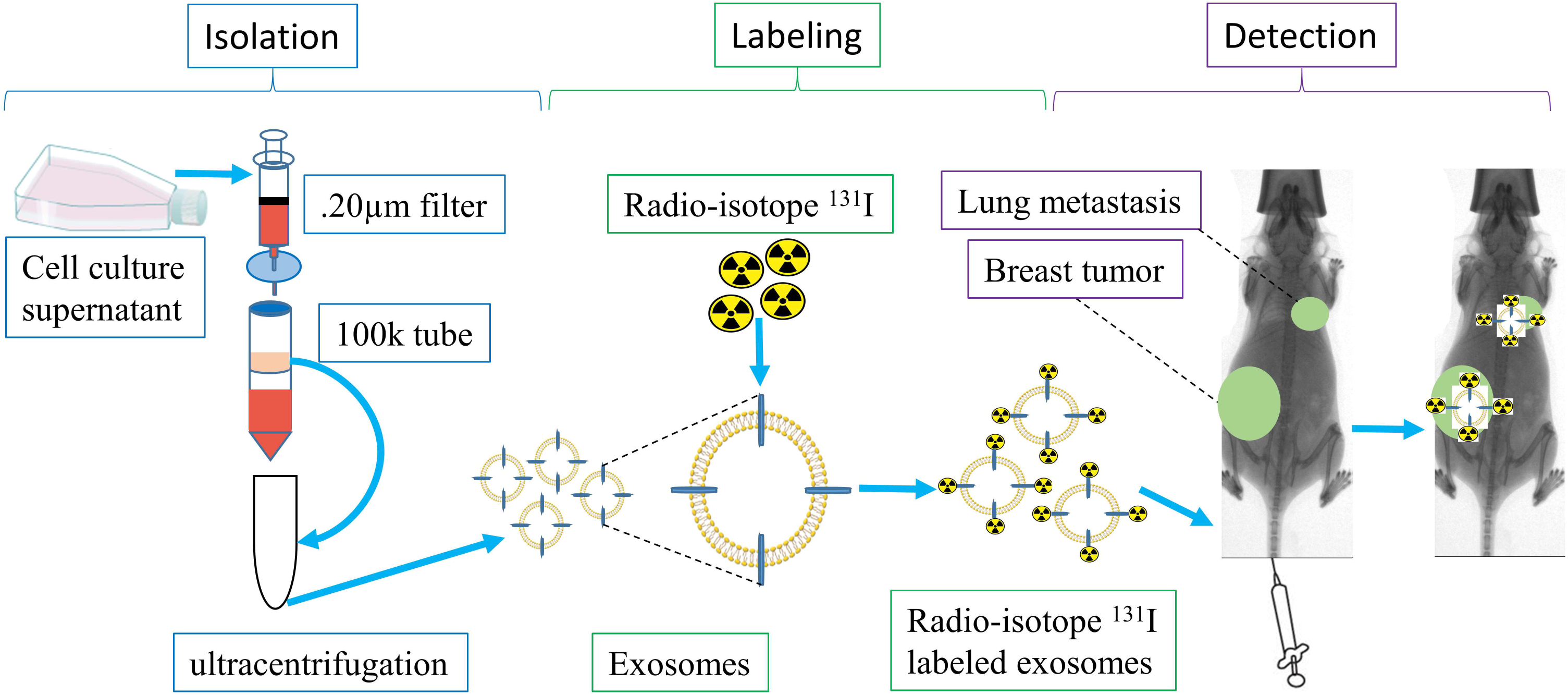

