## Supplemental materials for "Differential *in vivo* biodistribution of ^131^ I-labeled exosomes from diverse cellular origins and its implication in the theranostic application"

**Cell lines**

4T1, a murine mammary carcinoma cell was originally obtained from the American Type Tissue Culture Collection (ATCC), and modified by Dr. Hassan Korkaya (Augusta University) to express the luciferase gene reporter. Murine AT3 breast cancer cell line was obtained from Dr. Kebin Liu of Augusta University. For cell cultures and propagation, both cells were grown in Roswell Park Memorial Institute 1640 medium (RPMI) (Thermo Scientific), supplemented with 10% fetal bovine serum (FBS) (Nalgene-GIBCO), 2mM glutamine (GIBCO, Grand Island, NY, USA) and 100U/mL penicillin and streptomycin (GIBCO, Grand Island, NY, USA) at 5% CO_2_ at 37 °C in a humidified incubator. For the generation of exosomes, both cells (5x10^6^ cells in T175 flask) were grown in RPMI-1640 media containing 10% exosome free FBS and incubated in a humidified incubator in hypoxic condition (1% oxygen) for 48 hours. Human embryonic kidney 293 cell line (HEK293) was obtained from Dr. Satyanarayana Ande of Augusta University and was grown in Dulbecco's Modified Eagle Medium (DMEM) (Corning, NY, USA) containing 10% exosome free FBS.

**Nanoparticle tracking analysis**

Nanoparticle tracking analysis (NTA) was performed using ZetaView, a second-generation particle size instrument from Particle Metrix for individual exosome particle tracking as described preciously [26-28]. This is a high performance integrated instrument equipped with a cell channel, which is integrated into a ‘slide-in’ cassette and a 405-nm laser. Samples were diluted in 1X PBS between 1:100 and 1:2000 and injected in the sample chamber with sterile syringes (BD Discardit II, New Jersey, USA). All measurements were performed at 23°C and pH 7.4. As measurement mode, we used 11 positions with 2 cycles, and for analysis parameter, we used maximum pixel 200 and minimum 5. ZetaView 8.02.31 software and Camera 0.703 μm/px were used for capturing and analyzing the data.

**Flow cytometry**

The common exosome markers, mouse-specific anti-CD9 FITC, and anti-CD63 APC antibody (Biolegend, San Diego, CA, USA) were used to label exosomes at 4 °C for 30 minutes. Flow cytometry samples were acquired using Accuri C6 flow cytometer (BD Biosciences) with the threshold set at 10 and analyzed by BD Accuri C6 software.

**Isolation of myeloid-derived suppressor cells (MDSCs) and endothelial progenitor cells (EPCs)**

MDSCs were isolated from spleens of tumor-bearing mice 3 weeks after orthotopic tumor cell implantation using anti-Mouse Ly-6G, and Ly-6C antibody-conjugated magnetic beads (BD Biosciences, CA, USA). The purity of cell populations was > 99%. In short, the spleen was disrupted in PBS by using the plunger of a 3mL syringe, and then cell aggregates and debris were removed by passing cell suspension through a sterile 70μm mesh nylon strainer (Fisherbrand™). Mononuclear cells were separated by Lymphocyte separation medium (Corning®, NY, USA) as white buffy coat layer. Cells were then centrifuged at 1500 rpm for 10 minutes followed by a washing step with PBS at 1200 rpm for 8 minutes. Then cells were resuspended at 1x10^8^ cells/mL in PBS and antibody conjugated with magnetic beads were added followed by incubation at 4°C for 30 minutes. Finally, positive cells were collected using a MACS LS column (Miltenyi Biotec, Germany) and a MidiMACS™ magnetic stand followed by a wash step with an extra amount of PBS. The purity of isolated MDSCs was checked by flow cytometry using Gr1 FITC and CD11b APC antibodies (purchased from Biolegend). MDSCs were grown in exosomes depleted media consisting of RPMI, 2mM L-glutamine, 1% MEM non-essential amino acids, 1mM sodium pyruvate and 10% FBS, supplemented with 100 ng/mL of GM-CSF. EPCs were isolated by a similar technique from the bone marrow of normal mice using CD117 (c-kit) microbeads (Invitrogen™). The purity of the isolated EPCs was determined by flow cytometry using CD117 APC antibody. After isolation EPCs were grown in a exosomes free media consisting of stem cell media, 2mM L-glutamine, 20ng/mL stem cell factor (SCF), 25 ng/mL thrombopoietin (TPO), 20ng/mL FMS-like tyrosine kinase-3 (FLT3). Exosomes were isolated after 48 hours, following the similar technique used to isolate tumor-derived exosomes.

**Protein quantification**

Isolated exosomes resuspended in a minimal amount of PBS were lysed by RIPA buffer with protease and phosphatase inhibitor (100:1 dilution). Then exosomal protein was quantified by Bradford assay using Pierce™ BCA Protein Assay Kit (Thermo Scientific™) and serial dilution of BSA standard (Thermo Scientific™).

**Thin layer paper chromatography for radiolabeling efficacy and stability**

3MM Whatman® cellulose chromatography paper was cut into 1×8 cm small pieces. The bottom spotted point was made by 5µL of each sample followed by submerging the bottom part of each piece (below the spotted point) into the eluent consisting of 100% methanol and 2M Sodium acetate solution (1:1). Then the pieces were allowed to remain upright until the eluent reaches the top part. The pieces were cut into the top and bottom halves and were subsequently put in the glass tubes for the measurement of emitted gamma activity by Perkin-Elmer Packard Cobra II Auto-Gamma. Total radioactivity was calculated by combining the activity from top and bottom halves. To determine the percent dissociation of bound ^131^I from exosomes, labeled exosomes were challenged with serum at 37ºC up to 48hrs. At different time points, free ^131^I, and serum challenged labeled exosomes were tested using thin layer paper chromatography as described above to determine the percent of bound vs. free ^131^I.

**Quantitative analysis of radioactivity in individual organ**

Reconstructed analyze formatted file was used in ImageJ (Wayne Rasband, National Institutes of Health, USA) version 1.51a for both CT and SPECT analysis. The primary tumor, a metastatic site in the lungs and other organs were identified by orthogonal, dorsal and ventral views from the resliced stack images. Z stack images were created from the CT and SPECT of the individual organ for depth and anatomical accuracy of the organ. Total radioactivity was determined by the sum of the values of the pixels (RawIntDen) in the selected region of interest (ROIs) around the organs. The activity in the individual organ was expressed in percent of activity in the whole body (total radioactivity dose).

***Ex vivo* gamma activity**

After the final scan, animals were euthanized, and their organs were harvested and weighed. Emitted gamma radiation from each organ was measured by Perkin-Elmer Packard Cobra II Auto-Gamma after transferring them into the individual glass tube.

**Statistical analysis**

Quantitative data were expressed as mean ± standard error of the mean (SEM), and statistical differences between groups were determined by analysis of variance (ANOVA) followed by multiple comparisons using Tukey’s multiple comparisons test. GraphPad Prism version 7.03 for Windows (GraphPad Software, Inc., San Diego, CA) was used to perform the statistical analysis. Differences with p-values less than 0.05 were considered significant (*p<.05, **p<.01, ***p<.001, ****p<.0001).

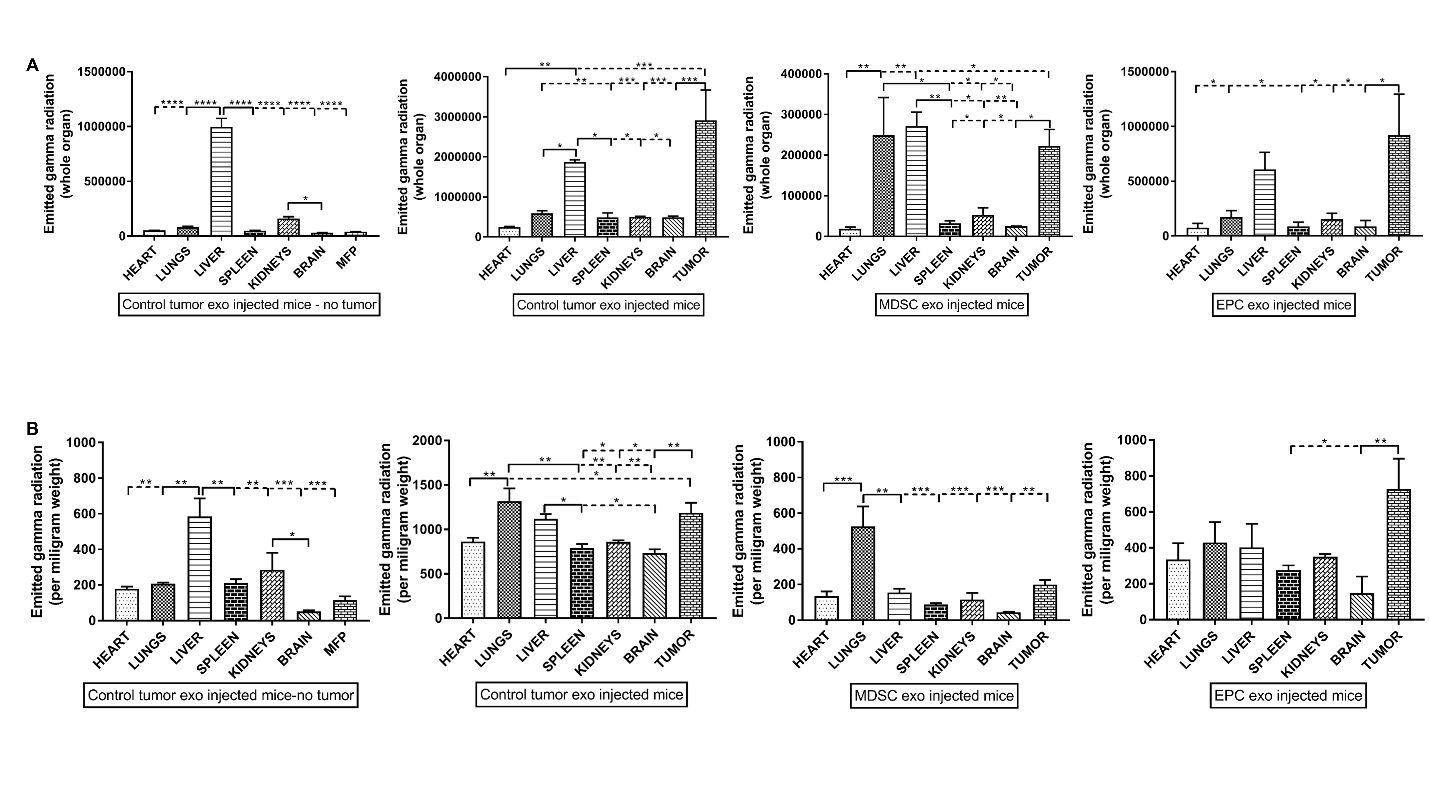

**Figure S1.**

*Ex vivo* quantification of radioactivity by gamma counter after final SPECT/CT scan followed by euthanasia and harvesting different organs. **(A)** As a whole organ-wise, only the tumor and liver retained noticeable radioactivity in all the groups. Lungs from the mice injected with MDSC exo showed significant radioactivity. **(B)** Quantification of radioactivity per milligram of the weight of an individual organ that reveals resembling changes of radioactivity as of the whole organ. Quantitative data are expressed in mean ± SEM. *P<.05, ***P<.001, ****P<.0001. n = 3.

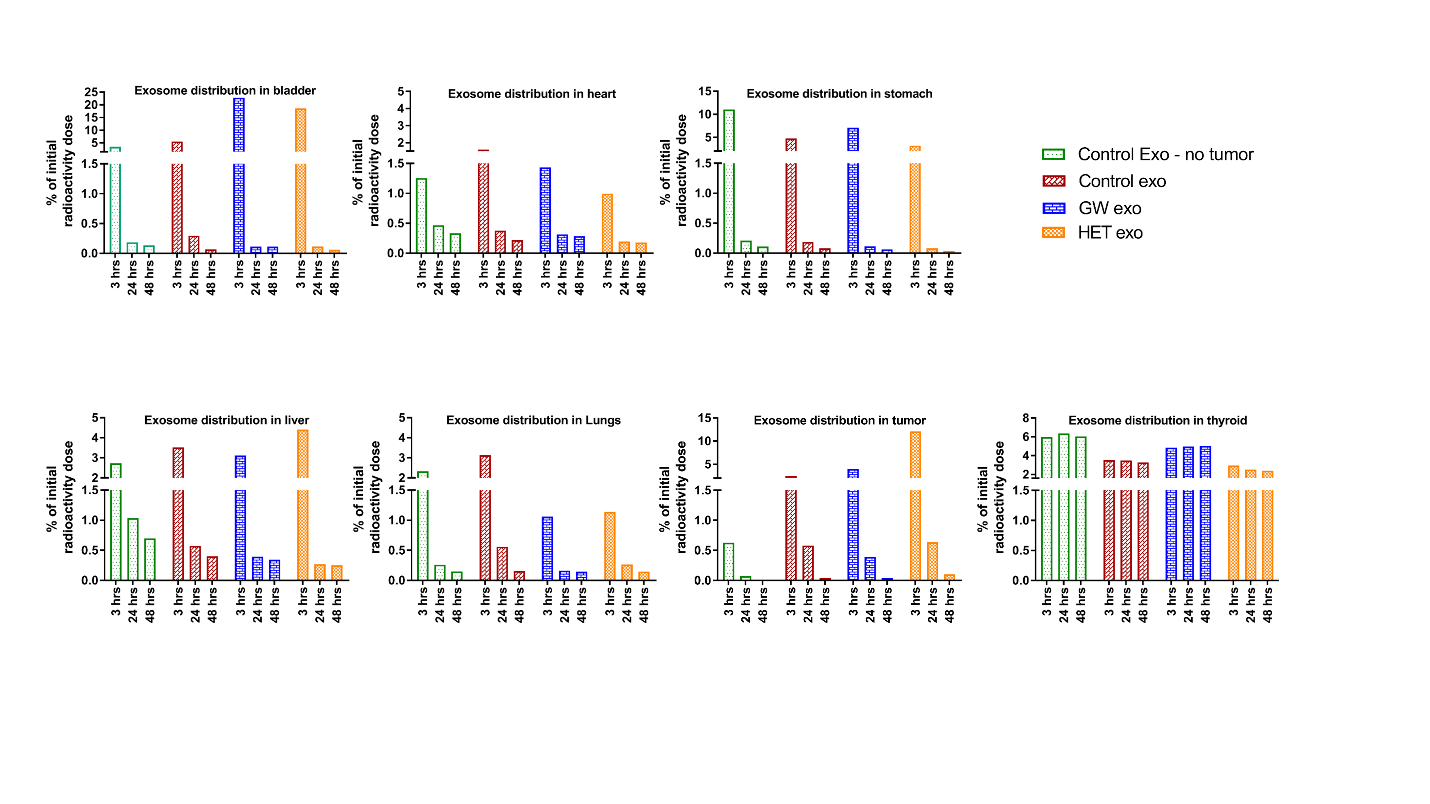
**Figure S2.**

Quantitative biodistribution of tumor-derived exosomes in tumor-bearing or non-bearing mice (representative from each group) for the bladder, heart, stomach, liver, lungs, tumor and thyroid after 3 hrs, 24 hrs and 48 hrs of intravenous injection of ^131^I labeled exosomes. During the first scan (3 hours), highest radioactivity was measured in the bladder, indicating one of the routes of excretion of ^131^I labeled exosomes. Albeit, most of the exosomes were cleared out of the body before the second scan (24hours), there was some radioactivity remained in the tumor, lungs, and liver in some groups. There is no apparent radioactivity difference in thyroid over the time, implying no dissociation of ^131^I from the labeled exosomes that would subsequently increase the activity in the thyroid.
